# A minimally invasive floating-wire interface for transcranial deep brain stimulation

**DOI:** 10.64898/2026.03.04.709293

**Authors:** Vishal Jain, Mats Forssell, Pulkit Grover, Maysam Chamanzar

## Abstract

**Background:** Non-invasive neuromodulation technologies have advanced considerably. Yet, precise and focal activation of deep brain regions remains challenging due to the rapid attenuation of electric fields across the scalp, skull and brain surface.

**Objective:** We present FLOATES (FLOAting Transcranial Electrical Stimulation), a novel approach that employs an untethered wire implanted in the brain which passively relays currents injected transcranially from the brain surface to deep brain regions, achieving focused stimulation deep within the brain.

**Methods:** We validated FLOATES through a combination of simulations, benchtop testing, and in vivo rodent studies. The benchtop experiments confirmed the ability to relay the field across the floating wire. Rodent studies demonstrated capability to stimulate deep brain regions in vivo.

**Results:** Our simulation and benchtop testing results indicate that FLOATES can deliver significantly higher electric fields to subcortical regions compared to conventional transcranial stimulation approaches. Further in-vivo results demonstrated deep subthalamic nuclei stimulation to evoke limb motor responses and demonstrated a significantly lower motor threshold compared to transcranial stimulation. Finite element simulations reveal that the efficiency of FLOATES depends on several key parameters including input field strength, wire length and diameter, exposed electrode area, impedance, and tip geometry. Simulations using a human-sized head model suggest that electric fields sufficient for brain stimulation can be obtained with reasonable currents injected to the scalp.

**Conclusion:** Together, these results establish a theoretical and experimental foundation for FLOATES as a minimally invasive and spatially precise brain stimulation platform in modulating deep neural circuits implicated in neuropsychiatric and movement disorders.

## 1. Introduction

Deep brain stimulation (DBS) has long been established as an effective treatment for various neurological disorders, including Parkinson’s disease, essential tremor, and dystonia [1]. Traditional DBS systems include electrodes that are surgically placed in specific brain regions, an electrical pulse generator and a wire that connects the pulse generator to the electrodes [2]. The pulse generator is usually implanted in the chest cavity and the wire is routed under the skin. While effective, the surgical procedure is highly invasive, with risks such as infection and hemorrhage [2, 3]. In addition, other post-surgical complications can also happen such as electrode/lead migration, pulse generator malfunction and skin erosion [4, 5]. Many of these complications happen because of the wired connection between the deep brain electrodes and the implanted pulse generator.

These risks have led to increased interest in non-invasive brain stimulation techniques targeting deeper regions in brain. Among these techniques, focused ultrasound stimulation (FUS) has shown considerable promise. FUS leverages high-intensity sound waves to target specific brain regions without the need for incisions, offering precise modulation of neural activity [6, 7], but its effectiveness in clinical DBS treatments is not well established [8]. Other traditional non-invasive brain stimulation techniques such as transcranial magnetic stimulation (TMS) or transcranial electrical stimulation (TES) offer promising alternatives to surgical methods but come with their own set of challenges and limitations. Most importantly, the depth and intensity of stimulation achievable by TMS and TES are limited compared to implanted electrodes, essentially restricting their use to surface or cortical brain regions rather than deeper structures involved in the neurological disorders treated by DBS [9, 10]. This is because the electric or magnetic fields enter the brain from the surface and have amplitudes that decay as they travel deeper in the brain. In order to achieve useful evoked response in the deeper regions, the field amplitudes in the shallower brain regions will necessarily be suprathreshold, resulting in side effects and likely severe safety concerns. In order to increase the depth that can be reached by TES, temporal interference (TI) utilizes two pairs of scalp electrodes to inject sinusoidal currents at slightly offset frequencies to produce neuromodulatory effects in deeper regions without superficial stimulation [11] [12], but the effectiveness of the technique is under debate [12], and no clinical trials for suprathreshold stimulation have been reported.

Another recent approach to reduce the surgical burden of DBS is the development of miniature stimulators that can be mounted in the skull and therefore do not require placing an implanted stimulator in the chest and connecting it to the DBS leads [13]. But the size of these devices requires a large craniotomy which still presents high surgical risks. Smaller stimulators are currently being developed [14] but have not been demonstrated over long-term or for DBS applications.

An alternative approach that has been employed in peripheral nerve stimulation (StimRouter, Bioness Inc.) consists of only implanting a passive lead consisting of a stimulation electrode positioned in close proximity to the target nerve at one end and a capture electrode shallowly placed under the scalp at the other end. Current pulses are injected using a wearable stimulator and couple transcutaneously to the implanted lead, resulting in stimulation of the nerve [15]. This technique has been used in clinical trials in multiple conditions targeting peripheral nerves with remarkable safety and efficacy results [15], but to date, no fully passive implant has been demonstrated for DBS. Deep brain stimulation using transcranial methods necessitates a new approach to achieve the required amplitude in deep brain regions while enhancing targeting precision, reducing individual variability, and improving overall safety.

In the current manuscript, we provide a novel method, called FLOATES (FLOAting Transcranial Electrical Stimulation), a minimally invasive technique that combines the benefits of DBS and TES to enable deep brain stimulation via transcranial electrodes. FLOATES consists of the surgical implantation of a free-floating wire in the brain, targeting the desired region (Figure 1). The wire has exposed electrodes at both ends, but is otherwise insulated. The distal (output) electrode acts similarly to a conventional DBS electrode in locally stimulating a targeted area, and the proximal (input) electrode is located at the top of the brain. Following implantation, electrodes placed on the scalp are used to non-invasively inject currents into the brain. We have recently demonstrated that focal and steerable electric stimulation can be delivered to the brain through skull non-invasively using high density patterns of electric stimulation on the surface [16]. The use of high-density scalp electrodes for stimulation allows designing focal currents that couple into the wire via the input wire electrode, allowing stimulation through the output electrode. This technique potentially reduces many of the hardware-related complications of DBS, while providing similar outcomes. Because the wire is not physically connected to the stimulation circuit, the surgical procedure is greatly simplified. Additionally, long-term risks of infection are also likely reduced due to less skull removal, and hence a likely complete recovery of the skull. Wearable scalp electrodes are connected to an external stimulator, which also avoids complications due to malfunction of the implanted hardware, or infection risks at the site of the implanted hardware [17]. In this paper, we first demonstrate this concept through simulations and measurements of the electric field generated by this technique, then report results of in vivo experiments in a mouse model showing functional effects as well as validation at the cellular level. We report parametric simulations of performance as a function of wire parameters for mouse- and human-sized head models.

**Figure 1.**
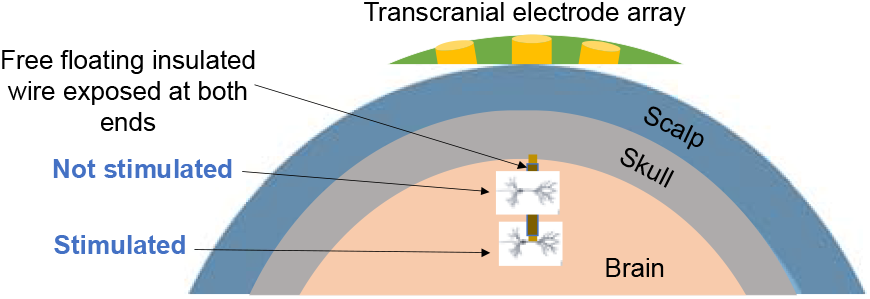
Schematic to represent FLOATES concept

**Figure 1.**
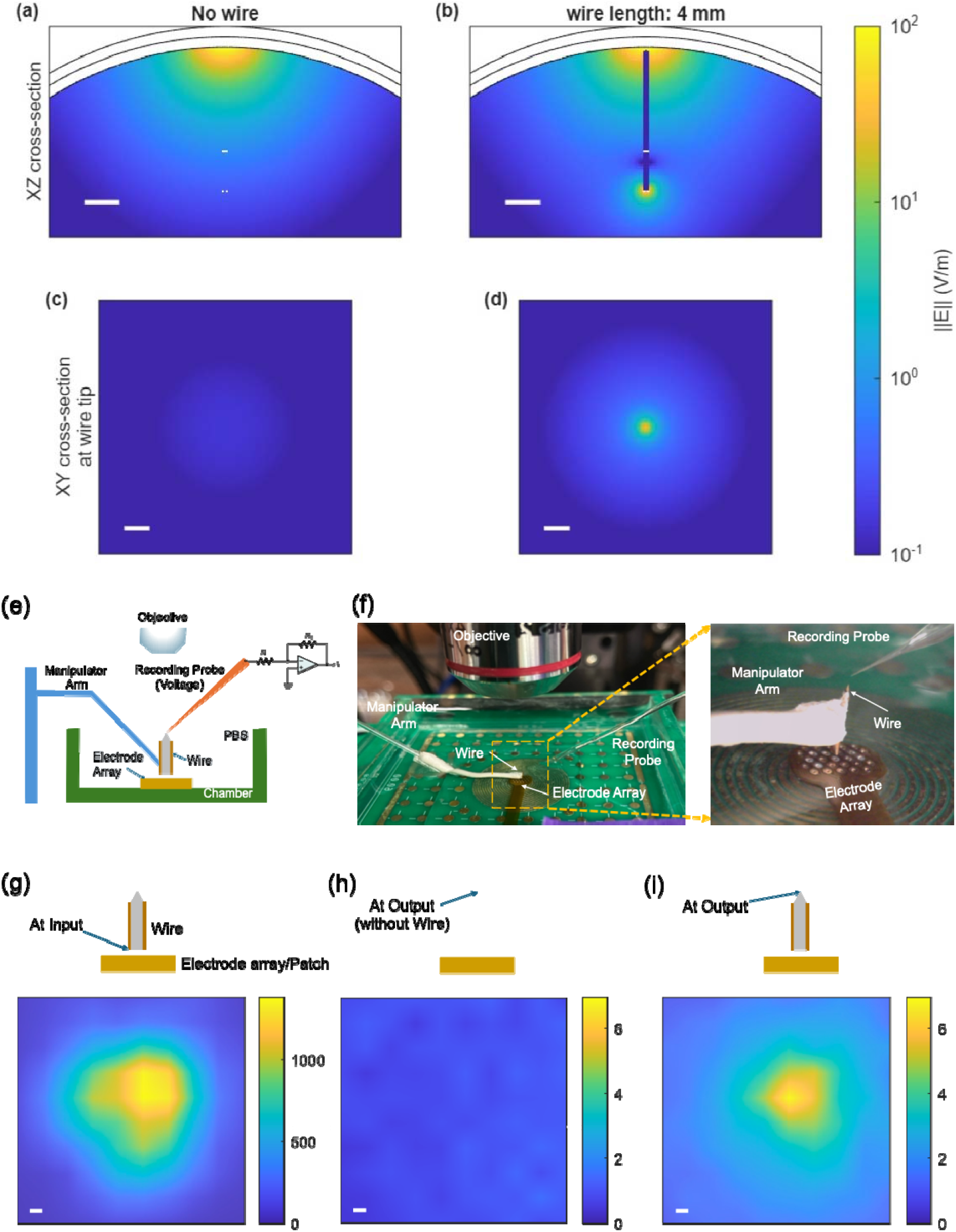
Simulation and benchtop demonstration of current coupling to deep regions. (a-d): Finite element simulation of electric field generated in the brain. (a,c): by a transcranial scalp electrode array and (b,d): relayed using a floating wire (diameter=200 µm, length=4 mm, scale bar: 1 mm). (e-i): Benchtop characterization of FLOATES. (e): schematic. (f): experimental setup. (g-i): Electric field magnitude (V/m) measured in saline, normalized to a 1 mA injected current, in three conditions: (g): at the location corresponding to the input of the wire (measured in the absence of the wire), 200 µm from the electrode array; (h): at the location of the wire output in the absence of the wire (scale bar: 100 µm); and (i): at the output of the wire (4.2 mm above the current-injection electrodes).

## 2. Methods

### 2.1. Tetherless wire preparation

To make tetherless wires, we used concentric metal microelectrodes (Model PI2CEA3-200; MicroProbes for Life Science, Gaithersburg, MD, USA) for benchtop and in vivo FLOATES experiments. Each wire was inspected under a stereomicroscope to confirm structural integrity and insulation continuity. To achieve different insertion depths suitable for bench and in vivo preparations, the wires were trimmed to lengths ranging from 3.5 mm to 4.0 mm using a wire cutter. Following trimming, the insulation was examined to verify that only the distal tip remained exposed, while the remainder of the shaft maintained intact insulation. Electrodes were cleaned in sterile saline and air-dried prior to use. These length-adjusted electrodes were subsequently used in both benchtop validation studies and in vivo experiments to achieve precise positioning within the target region.

### 2.2. Benchtop experiment

For benchtop testing, the tetherless wires were connected to a motorized manipulator arm and placed in a tub containing 1x phosphate buffered saline (PBS). The bottom of the tub contains a flexible printed circuit board (FPCB) consisting of an electrode array (ring electrodes with outer diameter: 0.35 mm and inner diameter: 0.15 mm, arranged in a hexagonal grid at a 0.65 mm pitch). In this work, we only use a single electrode as the anode and the six surrounding electrodes as the cathode. Using the micromanipulator, the floating wire was precisely aligned above the central anode, with the bottom of the wire 200 µm above the electrodes (corresponding to the thickness of a mouse skull). We apply pulsed current stimulation using biphasic, charge-balance 1 ms/phase pulses with 0.3-1.2 mA injected through the electrodes (the current was adjusted to ensure the recording amplifier was not saturated). We measured the voltage potential created in the PBS using a PBS-filled glass pipette connected to an amplifier and digitizer (Axon Instruments). The pipette recorded the potential field in a 100 µm grid. The electric field can then be calculated using:

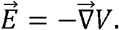

We recorded the electric field at a distance of 4.3 mm from the electrodes (corresponding to 100 µm from the tip of the wire) both with and without the floating wire. In the absence of the wire, we also measured the electric field 200 µm above the electrodes (corresponding to the location of the input of the wire).

### 2.3. Animals

For current study, C57Bl6 mice (12-18 weeks old) for either sex were used. All mice were randomly distributed among different groups. All experiments involving animals were performed in accordance with the Institutional Animal Care and Use Committee guidelines. The use of animals and all procedures were approved by Carnegie Mellon University’s Institutional Animal Care and Use Committee (IACUC). Animals were maintained on a 12 h light-dark cycle with free access to food and water.

### 2.4. Intracortical microstimulation (ICMS) mapping

Initial experiments using invasive probes were performed to identify the target region, i.e. the portion of the subthalamic nucleus (STN) that results in motor evoked potentials (MEPs) in the contralateral forelimb when stimulated electrically. C57 mice (n=2) were anesthetized using a ketamine/xylazine cocktail and headfixed on a stereotaxic frame. A microwire electrode (PI2PT30.01A5, Microprobes) was inserted at five locations surrounding the coordinates of the STN (AP=-2.7mm+-1mm, ML=1.8mm+-1mm). Electrical stimulation (Ripple Grapevine and micro2+stim headstage, Ripple Neuro) was performed using biphasic pulses of 1000 µs pulse width were injected in trains of 7 pulses at 350 Hz, similar to prior work on transcranial stimulation in motor cortex [16]. The injected current amplitude was increased until motor threshold was achieved. Electromyographic recordings were performed using monopolar needles inserted in the contralateral upper limb muscles, connected to a differential amplifier/digitizer (Intan differential headstage, Intan Technologies) recording at 30 kS/s. Motor threshold was characterized using the peak-to-peak amplitude of the MEP following stimulation, as the smallest current able to achieve a MEP amplitude above 50% of the maximum MEP amplitude recorded. At each insertion location, stimulation was delivered at six different depths (DV0.8mm, 1.5mm, 2mm, 3mm, 4mm, 5mm) and the threshold required to evoke MEPs in the contralateral forelimb was measured.

### 2.5. In-vivo stimulation

FLOATES was validated in C57 mice by targeting the STN in the deep brain and recording motor evoked potentials MEPs in upper limbs. Mice (n=4) were anesthetized using a ketamine/xylazine cocktail and head fixed on a stereotaxic frame. An electrode array (ring electrodes with outer diameter: 0.35 mm and inner diameter: 0.15 mm, arranged in a hexagonal grid at a 0.65 mm pitch), with a hole (0.4 mm diameter) at the center was placed on the skull, with the hole above the location of STN (AP=-2.7 mm, ML=1.8 mm from Bregma). The electrode array was connected to a multichannel stimulator (Ripple Grapevine and micro2+stim headstage, Ripple Neuro). A stimulation pattern utilizing 6 cathode and 6 anode electrodes surrounding the hole was used for stimulation. Stimulation waveform and EMG recording for motor threshold were performed as described in the previous section.

The motor threshold was characterized in three conditions, without moving the electrode array or the recording electrodes: (1) “intact skull” condition: transcranial stimulation through intact skull, without floating wire, as described above; (2) “skull with hole” condition: a hole (0.3 mm diameter) was drilled in the skull through the hole in the electrode array (i.e. at the identified STN location); and (3) “hole+wire” condition: a floating microwire (as above, consisting of the tip of a conical microwire (PI2CEA3-200, Microprobes) cut to a length of 4.5 mm) was inserted through the same hole in the skull, such that its proximal end is flush with the brain surface and its distal end reaches the STN.

### 2.6. Finite element simulation

Finite element simulations were performed using the electric currents interface in COMSOL Multiphysics. To mimic the conditions of rodent experiments, a spherical head model was utilized consisting of 3 layers: scalp (radius=10 mm, conductivity=0.465 S/m), skull (radius = 9.8 mm, conductivity =0.01 S/m), and brain (radius=9.6 mm, conductivity =0.2 S/m). Circular electrodes (0.4 mm diameter) were placed on the surface of the scalp, matching the arrangement of 7 electrodes used in the benchtop characterization (center electrode placed on the zenith, surrounded by six evenly distributed equidistant electrodes at a 0.65 mm distance). Electric current (1 mA) was injected into the center electrode, with the six surrounding electrodes acting as the ground (and therefore the return electrodes) and the electric field generated in the brain in steady state was simulated.

A cylindrical floating wire was included (nominal parameters: diameter=0.2 mm, depth from the brain surface=0 mm, length=4 mm, conductivity=10^6^ S/m). The wire is insulated along the length and exposed at the top (proximal end, or input facet) and bottom (distal end, output facet). Additionally, a portion of the length around the sides near the top and bottom was also exposed (nominal electrode length = 0.1 mm for top and bottom). These exposed regions correspond to the metal areas acting as electrodes through which current is coupled into and out of the wire. These electrodes are modeled as a surface impedance (nominal surface resistance: 1 Ω/m). Simulations were performed in the presence and absence of this wire, and also while sweeping the design parameters to evaluate the effect of wire properties on the electric field relay capability.

Three metrics were considered to characterize the floating wire performance: (1) the total electric current flowing through the wire; (2) the electric field amplitude at the output of the wire (maximum electric field 100 µm away from the wire); (3) the volume of tissue around the tip of the wire in which the electric field exceeds a chosen threshold of 100 V/m (approximating the stimulation volume).

### 2.7. Analytical model

A simplified analytical model was developed to predict the FLOATES outputs (current coupling and electric field at the output). The model assumes low coupling between the wire and electric field, and calculates the current through the wire based on the potential difference between the input and the output facets of the wire, assuming an existing electric field. Specifically, the model uses as its input an electric field potential in the brain produced in the absence of the floating wire (obtained from a finite element simulation or analytical expressions as shown in our previous work [18]. A cylindrical wire is defined by its position within this electric field, dimensions (radius *r* and length *L*), insulation coverage (quantified by the distance of deinsulated metal at the input 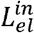 and output 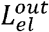 of the wire), bulk conductivity *ρ*_*wire*_, and surface resistance of the exposed metal *ρ*_*el*_. To calculate the current flowing through the wire,*V*_*in*_, the average potential along the exposed region at the input of the wire is calculated, and *V*_*out*_, the potential in the middle of the output face of the wire are first calculated. The total wire resistance *Z*_*tot*_ is the series combination of three resistances: *Z*_*in*_, the resistance of the input electrode; *Z*_*out*_, the resistance of the output electrode; and *Z*_*wire*_, the bulk resistance of wire:

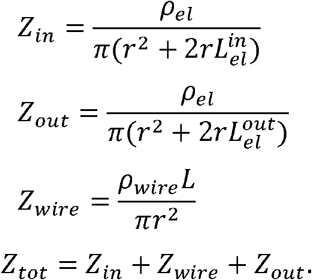

Additionally, the resistance path from the output of the wire (spreading resistance from the wire output) is approximated as:

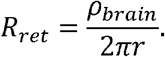

Then the current flowing through the wire is

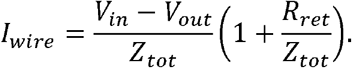

The electric field at the output of the wire is then calculated by assuming a constant current density along the entire exposed electrode at the output facet:

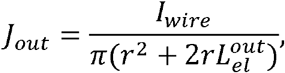

and the electric field at a point (x,y,z) is calculated based by integrating the distance between this point and the exposed metal electrode area of the output facet.

## 3. Results

### 3.1. Simulation and benchtop demonstration

Finite element simulations, illustrated in Figure 2(a-d), reveal that FLOATES can create significantly higher electric fields at deep brain regions compared to traditional transcranial electrical stimulation. The concept was also validated in the benchtop setting, as shown in Figure 2(e-f) using an electrode array placed at the bottom of a chamber filled with PBS. A floating wire was suspended in the solution 200 µm above the electrodes to mimic the thickness of a mouse skull. Figure 2(g-i) shows the electric field measured at the proximal (input) and distal end (output) of the wire. The results indicate 7X enhancement in the electric field intensity at the output when the wire is present (7 V/m), compared to the condition without the wire (1 V/m), measured at the same distance (4.2 mm from the electrodes) (Figure 2(h-i)). The measurements illustrate a localized field around the wire output, demonstrating the effectiveness of the wire in maintaining consistent field strength around the output region. Without the wire, however, only a weak diffuse field can be observed due to the distance from the electrodes, highlighting the wire’s role in conducting current to deeper regions.

**Figure 2.**
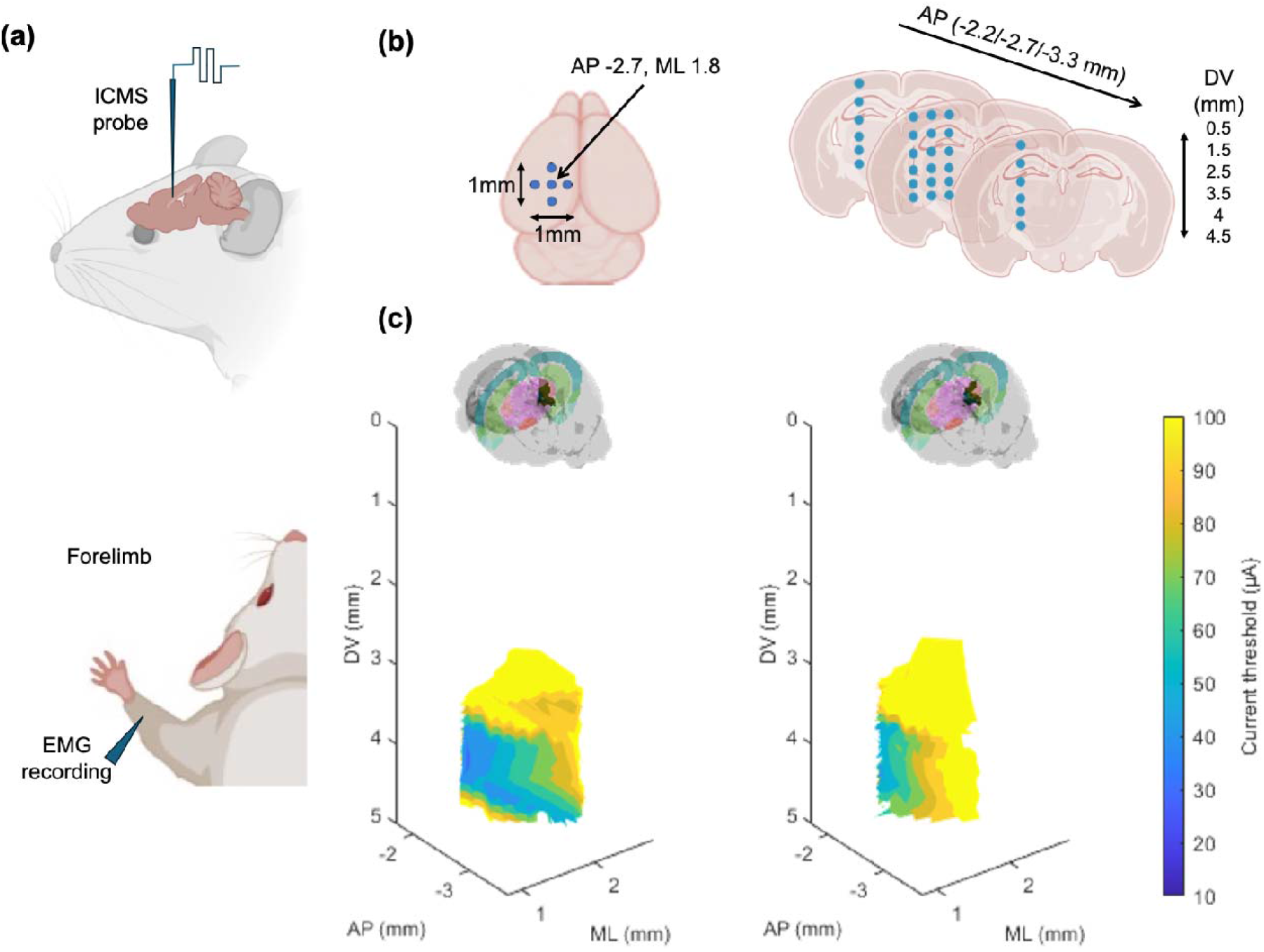
(a) ICMS mapping was performed targeting different locations and MEP response was recorded from contralateral forelimb.(b) ICMS probe insertion location at coordinates targeting STN and additional 4 nearby regions at step size of 0.5 mm in ML and AP and 0.5/1mm in DV. (b) Motor threshold (minimum current required to elicit MEPs in the region of interest. No motor response was observed at locations shallower than DV=3 mm. The overall responding location to evoke MEP in deep brain is very confined within 1mm^3^. Part of the figure (a-b) was created in BioRender. Chamanzar, M. (2025) https://BioRender.com/15z0uq5

### 3.2. Identification of target brain region

For in vivo demonstration of FLOATES, a challenge is to identify deep stimulation targets to insert the floating wire to elicit motor-evoked potentials (MEP). Previous reports have shown MEP responses can be elicited by stimulating the subthalamic nucleus (STN) [19] in the deep brain (AP-2.7mm, ML1.8mm, DV4.5mm). Because the transcranial currents that couple into the wire can also affect superficial cortical regions, the wire insertion location must be such that stimulation of the superficial regions at the insertion location does not cause motor activity. This is critical, especially since we have previously demonstrated non-invasive stimulation of motor cortex in mice using surface electrodes to evoke MEPs in forelimbs [16]. To avoid motor cortex stimulation, the chosen region for insertion (AP=-2.7 mm, ML=1.8 mm) corresponds to the secondary visual cortex (V2). A baseline intracortical micro-stimulation (ICMS) experiment was conducted in which a microwire stimulation probe was inserted at 5 different locations surrounding the coordinates of STN i.e., AP -2.7mm+-1mm, ML 1.8mm+-1mm (Figure 3a). At each insertion location, stimulation was delivered at six different depths (DV 0.8mm, 2mm, 3mm, 4mm, 4.5mm, 5mm) and the threshold required to evoke MEPs in the contralateral forelimb was measured (Figure 3b).

**Figure 3.**
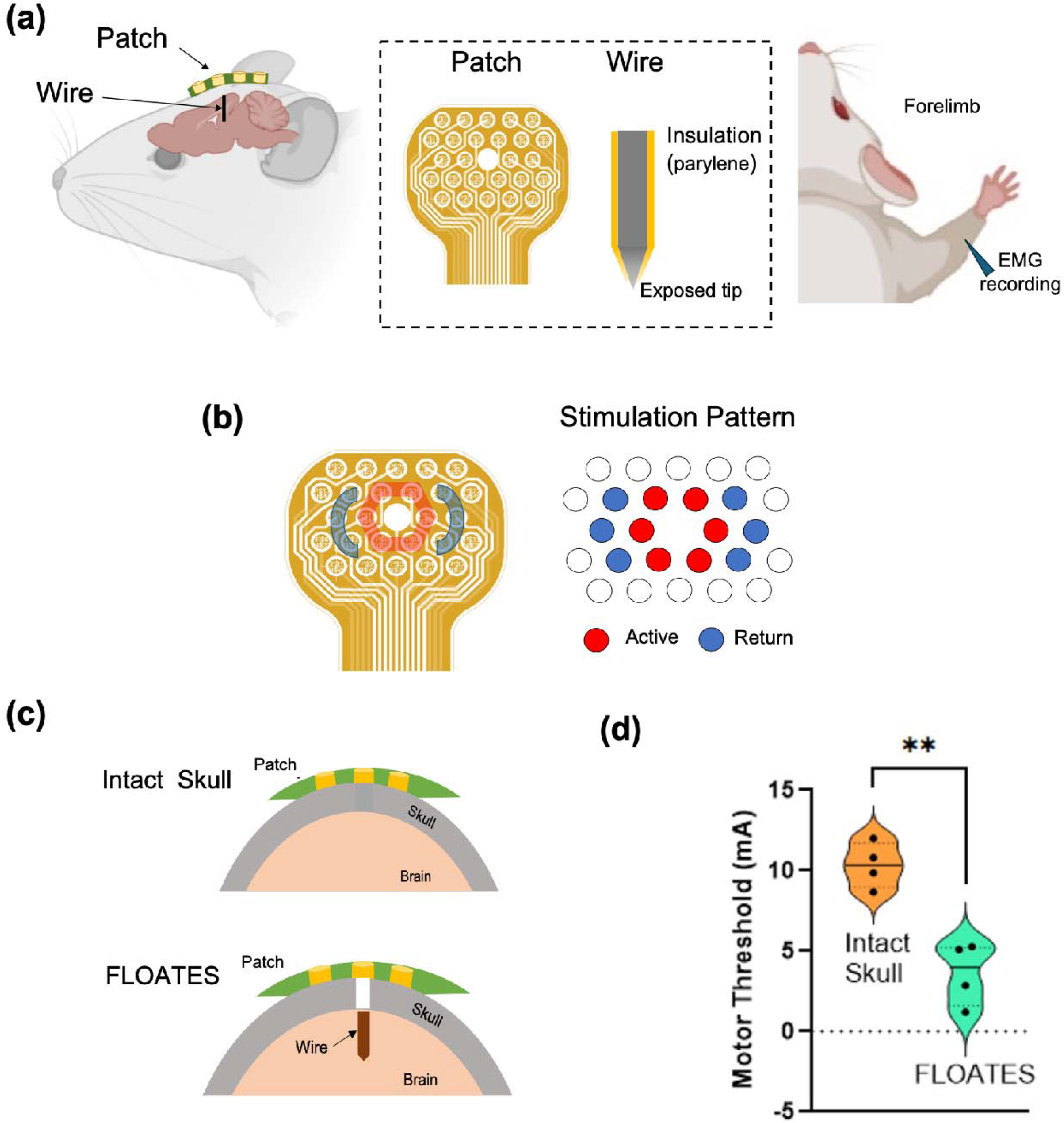
(a) Experimental setup in which the stimulation patch was placed on skull and a wire was inserted to target STN and motor response (MEPs) was recorded in contralateral forelimb. Created in BioRender. Chamanzar, M. (2025) https://BioRender.com/kt1hc00 (b) shows current injection pattern showing on the patch where circular arrangement of anode (red) electrodes, surrounded by cathode (blue) electrodes arrangement on both sides represented in the schematic as well on right.(c) shows the two experimental conditions: intact skull and FLOATES. (d) Motor threshold is plotted for each condition. Black dots indicate individual data points (n = 4). Dark and dashed lines represent median and percentile respectively. The shaded violin width reflects the kernel density estimate of data distribution. Statistical analysis was performed using unpaired t test with Welch’s correction and significance between groups is indicated by horizontal bars and asterisks (** p < 0.01).

Results revealed that the extent of the deep brain region in which stimulation can evoke motor responses spreads in AP and ML by 0.5 mm and in DV by 1mm (Figure 3b). The region is confined to depths greater than 4 mm, as no motor response was observed by stimulating more superficial layers (i.e. cortical and thalamic regions) (Figure 3c). This location was used for all subsequent experiments.

### 3.3. Stimulation of deep brain regions (STN) using FLOATES

FLOATES was validated in a mouse model by targeting STN in the deep brain and recording MEPs. Transcranial currents were injected into the brain using an electrode array placed on the mouse skull, and MEPs were recorded using EMG needle electrode inserted in the contralateral forelimb (Figure 4a). A floating wire was inserted into the deep brain targeting STN to demonstrate neuronal stimulation deep within the brain (Figure 4b). In each animal, two different conditions were tested: 1) transcranial stimulation without floating wire and an intact skull, and 2) transcranial stimulation with FLOATES wire inserted through the same hole in the skull. Another condition with just a hole in skull (with stimulation patch placed and no wire inserted) was performed to verify that the presence of the hole in the skull does not change the motor thresholds due to transcranial stimulation (Supplementary figure 1).

Results (n=4) revealed that FLOATES reduced the motor threshold by 3x (3.6 mA ± 0.97 mA; Mean ± SEM) when compared to stimulation through intact skull (10.33 mA ± 0.70 mA) with significance of p=0.0019 (Figure 4d). The condition with stimulation with a hole in the skull but no wire implanted, reduced the motor threshold but failed to achieve a statistically significant difference (threshold 7.3 mA ± 1.2 mA) (Figure S1).

### 3.4. Simulation study of FLOATES wire parameters

Our benchtop testing and in vivo animal experiments establish the feasibility of relaying a focused electric field current through FLOATES wires. As illustrated in Figure 2b, FLOATES can deliver significantly higher electric field amplitudes to deep brain regions compared to traditional noninvasive electrical stimulation. The rich design space of the FLOATES system enables optimizing the electric current coupling efficiency and the relaying mechanism. To understand the effects of different parameters on the performance of this system and inform future design and optimization strategies, we have performed rigorous

Finite Element Method (FEM) simulations to study the effect of the input field, the wire dimensions (length and diameter), the area of exposed electrodes, the electrode impedance and the tip geometry. We considered three output metrics of (i) electric current amplitude flowing through the wire, (ii) the electric field amplitude at the output of the wire, and (iii) the volume of tissue in which the local electric field exceeds a chosen threshold (approximating the stimulation volume). In addition to the FEM simulation, an analytical model was developed which decouples the wire from the electric field (i.e., it assumes that the presence of the wire does not affect the electric field generated by the stimulation electrodes). In addition to providing insight into the interactions between the wire and the electric field, this analytical model, due to its simplicity, can be used for optimization studies.

#### 3.4.1. Effect of the input field

Figure 5a shows that for a nominal wire geometry, the current coupled through the wire as a function of the local field at the distal end of the wire (i.e. at the input electrode) calculated in the absence of the wire. The plot shows a good linear fit, meaning that, for a given wire, the current that flows through the wire is proportional to the field at the location corresponding to the input electrode. Therefore, the optimization of the wire parameters and the currents injected through the scalp are decoupled. Regardless of the wire properties, the current coupled into the wire will be maximized if we maximize the electric field at the input facet. This can be explained by thinning of the wire to be a small perturbation to the applied transcranial electric field and the current coupled into the wire itself is small. If a large current couples to the wire, then the wire perturbs the applied electric field more strongly, and the effects of the wire on the electric field cannot be decoupled. From the perspective of deep brain neural stimulation, in addition to coupling to the wire, the applied electric field results in off-target superficial stimulation. Therefore, the optimization of the currents injected through the scalp (which elicit the electric field in the brain) should take into account both the amplitude of the electric field at the wire input (to maximize the current flowing into the wire) and the electric field at other superficial locations. In practice, a confined input field is desirable to reduce off-target effects, but this comes at the cost of reduced maximum amplitude [20]. Effects of the wire geometry on the output quantities are detailed in Figure 5b.

**Figure 5.**
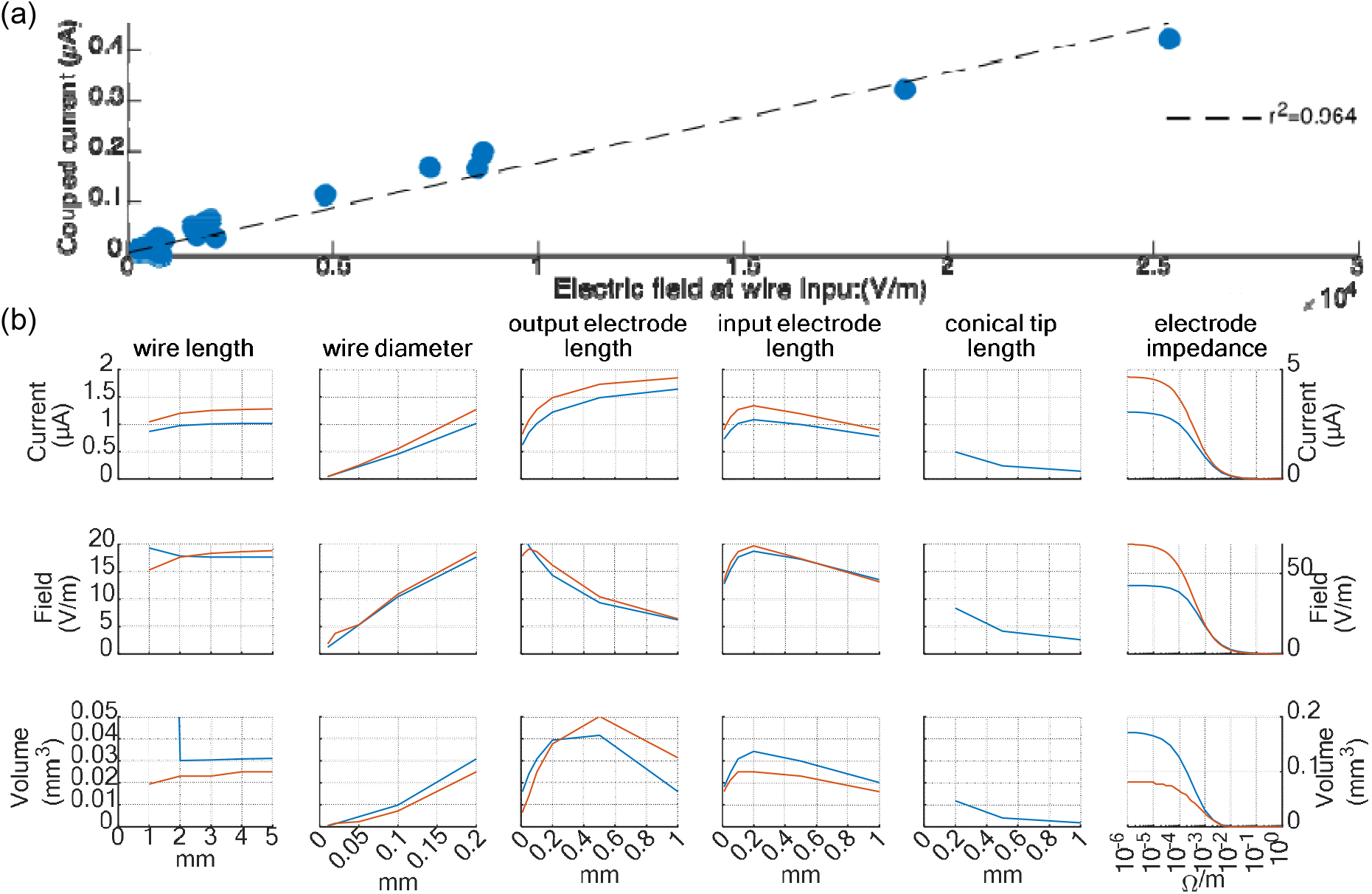
Simulations of FLOATES parameters on output quantities, assuming 1 mA current injected through the scalp. (a) Effect of the electric field at the input on the current flowing through the wire, for a fixed wire geometry. (b) Effect of wire parameters on (top) current coupling to the wire, (middle): electric field at the output, (bottom): volume above 100 V/m at the output. Blue: Finite element model; Red: analytical model. Plots in the same row are on the same y scale except for the right column

#### 3.4.2. Effect of the wire length

Both the current coupled to the wire and the output field indicate that unless the wire is very short (≤1mm), the output is not affected by the length of the wire. This is expected due to the high conductivity of the metal wire in the simulation, the conductivity of the wire is 10^6^ S/m, compared to 0.2 S/m for the surrounding tissue). For short wire lengths, the electric field at the output is larger because the effect of the applied electric field (resulting from the injected current through the scalp, without coupling to the wire) is still noticeable at the output.

#### 3.4.3. Effect of the wire diameter

Increasing the wire diameter reduces the wire impedance, and thereby increases the current coupled into the wire. While the current flowing into the wire is proportional to the diameter, the output field increases very little. This is because the output field is also inversely proportional to the area of the output electrode (which scales quadratically with the diameter).

#### 3.4.4. Effect of Exposed Area

We further examine the impact of varying the exposed area of the electrodes at both the input and output ends. For a cylindrical wire, increasing the exposed electrode area is achieved by increasing the length of the de-insulated portion of the wire. For the output electrode, increasing the exposed area reduces the overall wire impedance and therefore increases the coupling of current into the wire. However, the electric field at the output is inversely proportional to the exposed area; therefore, overall increasing the area reduces the electric field at the output. For the input electrode, increasing the exposed area also reduces the impedance, and therefore allows more current to couple into the wire. However, the benefits are limited because with larger electrode area, the average electrode depth also increases (since the area is increased by exposing a longer portion of the wire). Because the applied electric field decays with depth, the result is that a lower average electric field couples into the wire. Therefore, there is an optimal exposed input area to maximize the current coupled into the wire.

#### 3.4.5. Effect of Geometry at Output on Current Density at Output

Utilizing a conical tip, rather than a perfect cylinder reduces the current coupling and output field because the electrode area is reduced, thereby increasing the impedance of the wire.

#### 3.4.6. Electrode impedance

The electrode impedance is modeled as a surface resistance on the exposed metal area and is therefore proportional to the surface resistance *ρ*_*el*_. The plot shows that coupled current, output field and stimulation volume all decrease with increasing impedance (Fig. 6f). In practice for a given electrode size the impedance can be controlled in two ways: by changing the electrode material or depositing a surface coating (e.g. PEDOT) to reduce the electrode-tissue impedance, or by changing the injected signal through the electrode. The simulations are performed using steady-state models, and therefore, do not take into account the temporal properties of the signal. However, real microelectrodes behave like a capacitor, therefore their impedance is inversely proportional to the frequency of the signal. Using shorter stimulation pulses increases the characteristic frequency, and therefore, reduces the effective impedance.

**Figure 6.**
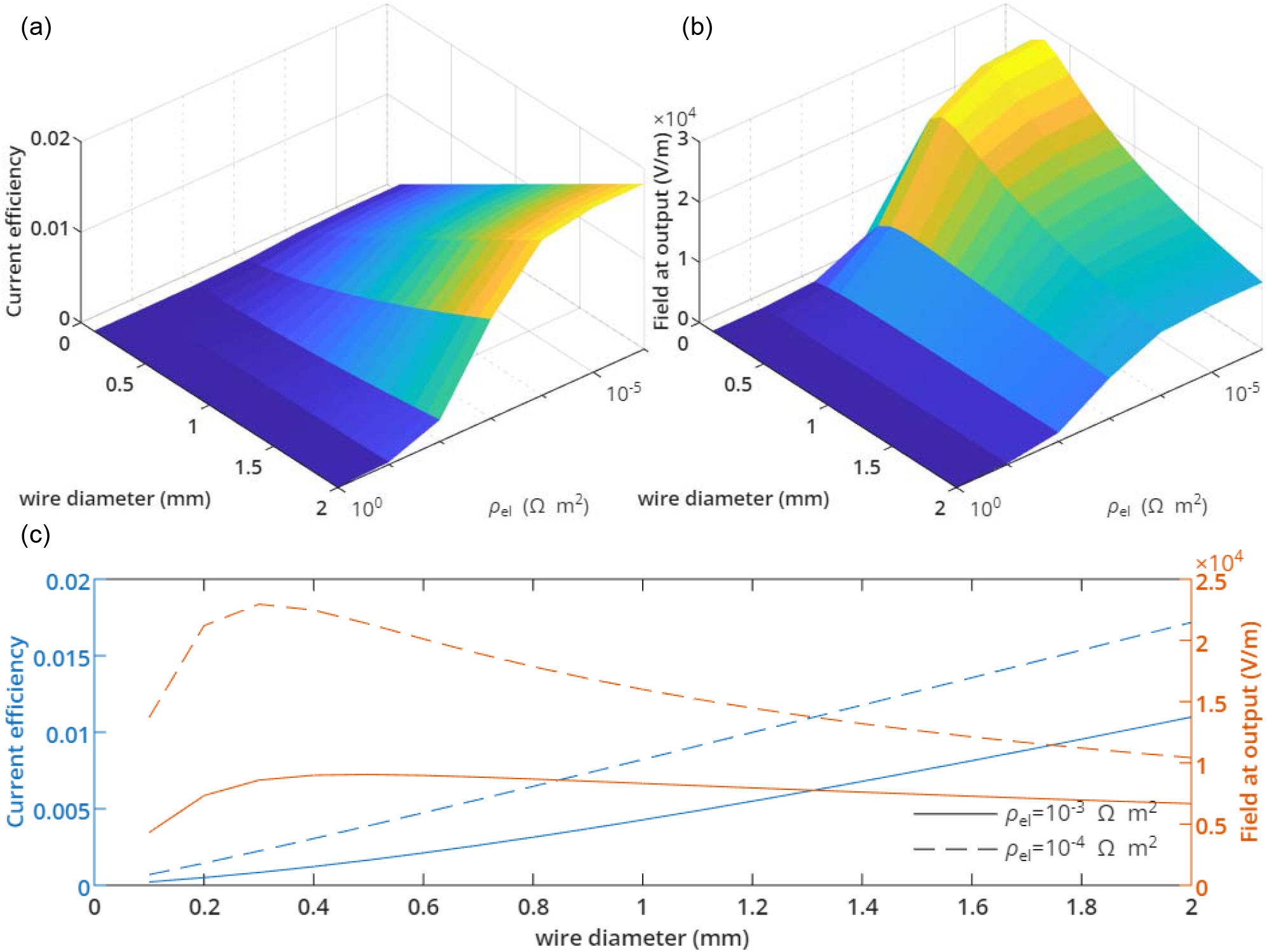
Analytical model simulations of FLOATES performance for a human-scale head (4-layer spherical head from [cite], R_head_=9.2 cm, t_csf_=1 mm, t_skull_=5 mm, t_scalp_=6 mm, with σ_brain_=σ_scalp_=0.2 S/m, σ_csf_=1 S/m, and σ_skull_=0.005 S/m). The floating wire is located at the surface of the brain (1.2 cm below the top of the scalp) and is 5 cm in length. The input transcranial field is generated by a “1+6” pattern (1 anode at the center, 6 cathodes symmetrically around it) with 7 cm anode-cathode distances. Simulations are performed using 1 A injected current. (a): current flowing through the wires (in A); (b): electric field at 100 µm from the wire; (c) line plots from (a,b) for two values of ρ_el_.

### 3.5. Simulation of coupling at human scale

To evaluate the feasibility of translating this approach to human-scale anatomy, we performed additional simulations using a simplified human head model. Fig. 6 shows the results of simulations performed using a human-scale head (4-layer spherical head from [21], R_head_=9.2 cm, t_csf_=1 mm, t_skull_=5 mm, t_scalp_=6 mm, with σ_brain_=σ_scalp_=0.2 S/m, σ_csf_=1 S/m, and σ_skull_=0.005 S/m) using a 5 cm–long wire with one end located at the surface of the brain. We swept the two most important wire parameters, the wire diameter and the electrode impedance and calculated the current flowing through the wire and the electric field at the output. Fig. 6c shows the outputs using electrode impedance in the order of magnitude of the wire used in experiments. Efficiencies of 0.4% to 0.8% are obtained with a 1 mm–diameter wire (similar to diameter of human DBS leads), meaning that 1A of current at the scalp results in 4-8 mA at the output of the wire. Realistic current that can be applied to the scalp using this electrode configuration is around 10-100 mA. When injecting 10 mA through the scalp, the electric field at the output of a 1 mm–diameter wire is 80-160 V/m, which is sufficient for neuronal activation. Using a more optimized 0.3 mm–diameter wire, fields of 230 V/m can be achieved with the lower impedance assumption (ρ_el_=10^-4^ Ω/m^2^, corresponding Z≈600 Ω).

## 4. Discussion

Noninvasive brain stimulation methods, such as transcranial magnetic stimulation (TMS) and transcranial electric stimulation (TES) have shown promise in targeting superficial cortical regions of the brain, but their effectiveness diminishes for deeper structures due to significant attenuation of fields as a function of distance [22] [23]. The findings from our benchtop experiments, simulations, and in vivo trials demonstrate the potential of FLOATES to deliver targeted stimulation to deep brain structures, by relaying surface-injected currents to deep brain regions using a floating wire.

Unlike traditional DBS that relies on a tethered connection between the implanted lead and a pulse generator, FLOATES provides effective deep brain stimulation with a far lower level of invasiveness, making it a suitable alternative in cases where minimizing surgical intervention and post-surgery risks is preferred. FLOATES uses a high-density surface electrode array for current injection, allowing it to steer currents/electric field to target the input facet of the floating wire with high accuracy.

Our simulations demonstrated that the current coupled into the wire is proportional to the input electric field, confirming that optimizing the wire design and input field strength can be decoupled for better control over the stimulation field at the distal end of the floating wire by enhancing the focal field at the input facet [23, 24]. However, care must be taken to avoid excessive off-target stimulation, a challenge commonly faced in non-invasive brain stimulation techniques like transcranial electrical stimulation (TES) [22, 25].

For the wire, the major factor influencing efficiency is the overall wire impedance, which is dominated by the input and output electrode electrochemical impedances. The electrode impedances can be reduced by increasing the exposed metal area, which for a cylindrical wire is achieved either through increasing the wire diameter or by increasing the length of the exposed area along the wire. For the input electrode, FLOATES could employ a wide conductive electrode resting on the brain surface, increasing the coupling efficiency as well as simplifying wire explantation. However, the output electrode impedance will then dominate the overall wire impedance. Reducing the output electrode impedance through geometric means (i.e. increasing the exposed metal area of the output electrode) will increase the current flowing through the wire but result in lower electric field at the output (because the amplitude of the electric field at the output of a wire is inversely proportional to the electrode area). Reducing the impedance can also be achieved through improved electrode-tissue interface (e.g. through electrodeposition of high surface area conductive materials such as PEDOT), without changing the geometry of the electrodes.

Our benchtop experiments were performed in homogeneous conductive media and at fixed electrode-tissue distances. While these rigorous experiments fully characterize the behavior of the FLOATES technology, they don’t capture the anisotropic conductivity, and field distortions that occur in vivo. Our in vivo mouse experiments, however, show that this method indeed works in real tissue to stimulate deep brain regions to the level that it can evoke downstream MEP responses in the limbs. While these responses confirm that stimulation propagates through the relevant motor circuits, quantifying and characterizing the stimulation zone at the distal end of the floating wire requires further experiments. Our in vivo experimental results in this paper show the efficacy and functionality of FLOATES. Demonstrating the chronic advantage of FLOATES over traditional DBS for long-term deep brain stimulation requires follow on studies. Finally, scaling this method to the larger brains of non-human primates and humans should be feasible by extending the length of the conductive floating wire. However, thicker scalp and skull layers would require redesigning the surface electrode array and current injection patterns. The larger head and brain size are more forgiving, allowing the use of larger surface arrays and thicker wires to mitigate current dispersion over longer distances.

In conclusion, these results support the feasibility of using a floating conductive wire to relay transcranially applied currents toward deeper brain regions with improved efficiency relative to surface stimulation alone, as demonstrated in rodent brain experiments and human head model simulations. While further studies are needed to characterize stimulation volumes, long-term stability, and performance in larger brains, this work provides a proof-of-concept for a minimally invasive strategy that could complement existing neuromodulation approaches by enabling deeper and more targeted field delivery without fully implanted systems.

---

**Supplementary Figure 1.**
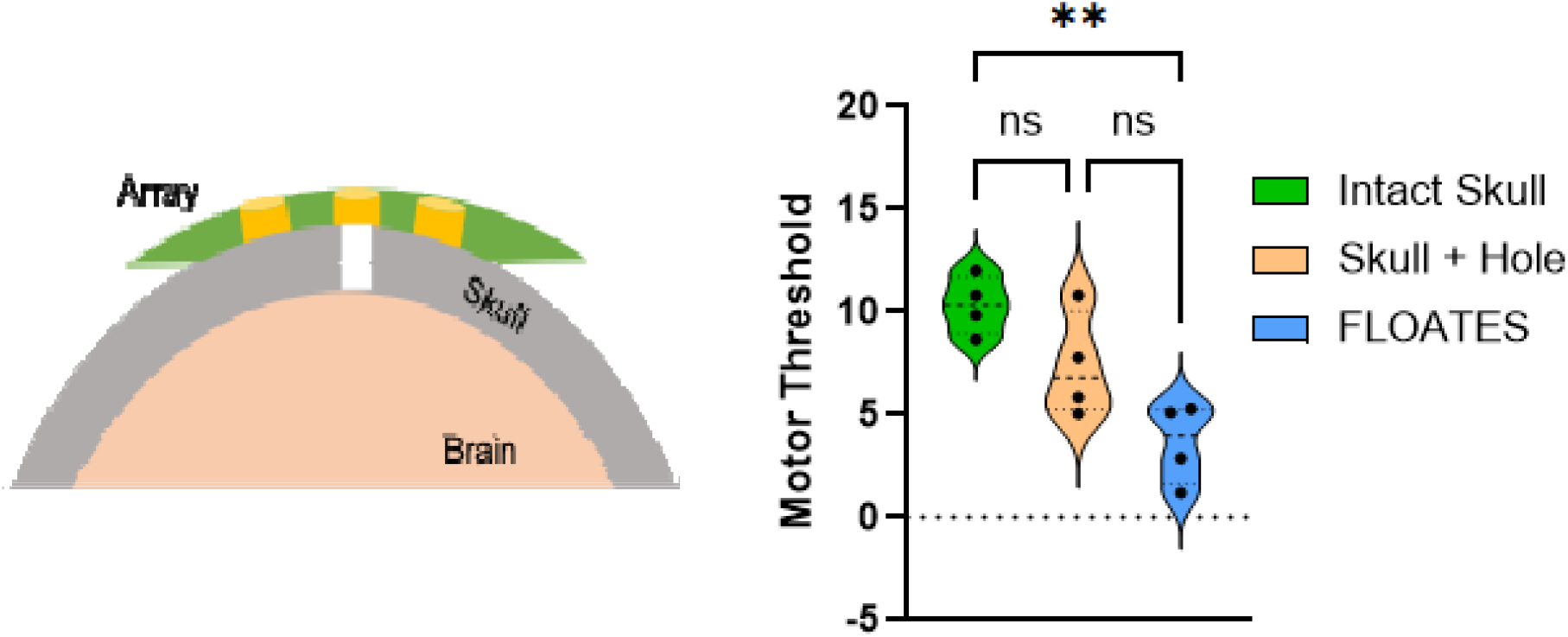
Data from Fig. 4d with the added group consisting of stimulation from a patch placed on the skull through which a hole has been drilled at the center (prior to insertion of the FLOATES wire). Motor threshold is plotted for each condition (including the other groups i.e. intact skull and FLOATES). Black dots indicate individual data points (n = 4). Dark and dashed lines represent mean and percentile respectively. The shaded violin width reflects the kernel density estimate of data distribution. Statistical analysis was performed using one-way ANOVA with Bonferroni post hoc test and significance between groups is indicated by horizontal bars and asterisks (** p < 0.01).

